# *Solanum americanum Bs2* and *ZAR1* homologs recognize *Xanthomonas euvesicatoria* effectors essential for infection

**DOI:** 10.64898/2026.07.27.741084

**Authors:** YoungWoo Koh, Hojin Jo, Jeongrae Kim, Haeun Cho, Injae Kim, Wanhui Kim, Chul Min Kim, Kee Hoon Sohn, Cécile Segonzac

**Author notes:** Corresponding author: C. Segonzac.

## Abstract

Multiple recognition events of pathogen-secreted effectors by immune receptors confer robust disease resistance in plants. Understanding the underlying mechanisms facilitates the discovery and deployment of valuable resistance genes for crop protection. *Xanthomonas euvesicatoria* causes devastating bacterial spot disease in solanaceous crops but cannot infect the wild relative *Solanum americanum*. Here, we identified *X. euvesicatoria* type III effectors (T3Es) that induce cell death in *S. americanum* when transiently expressed in leaf. By quantifying immune responses of the *S. americanum* SP2273 accession to *X. euvesicatoria* multiple-T3E knockout mutants, we demonstrated that at least nine T3Es (AvrBs2, XopAP, XopAU, XopE1, XopJ3, XopM, XopN, XopX, and XopZ1) collectively contribute to effector-triggered immunity (ETI). Among these, AvrBs2 and XopJ3 were the primary drivers of ETI, eliciting robust cell death and defense gene expression when naturally delivered into plant cells. We next generated *S. americanum* lines concomitantly edited at the corresponding immune receptor loci, *SaBs2* and *SaZAR1* (SP2273-*bz*). Genetic complementation of SP2273-*bz* confirmed that AvrBs2 and XopJ3 are specifically recognized by each of the four *SaBs2* homologs and by *SaZAR1*, respectively. Moreover, enhanced growth of *X. euvesicatoria* on the characterized SP2273-*bz* line indicated that both effectors are required for bacterial multiplication in *S. americanum*. Together, our findings provide a framework for understanding the mechanisms of ETI-mediated resistance and establish a genetic foundation for resistance breeding in solanaceous crops.

## Introduction

Leaf bacterial spot disease impacts tomato and pepper yield worldwide (Stall et al., 2009). This disease causal agent comprises distinct *Xanthomonas* lineages belonging to three species: *X. euvesicatoria*, *X. hortum* and *X. vesicatoria* (Osdaghi et al., 2021). The pathogenicity of these *Xanthomonas* species depends on the type three secretion system (T3SS) that delivers effector proteins into host cells to promote bacterial infection (Portaliou et al., 2015; Xin et al., 2018). Extensive genomic analyses and machine learning-based approaches identified at least 38 type III secreted effectors (T3Es) in the reference *X. euvesicatoria* strain *Xe* 85-10 (Schulze et al., 2012; Thieme et al., 2005; Teper et al., 2016; White et al., 2009). T3E families can be conserved across evolutionary distant bacterial pathogens, such as the YopJ effector family, represented in animal (*Yersinia enterocolitica*, *Salmonella enterica*, *Vibrio parahaemolyticus*) and plant (*Pseudomonas syringae*, *Ralstonia solanacearum*, *Erwinia amylovora*, *Acidovorax citrulli*, *Xanthomonas* spp.) pathogens reflecting essential roles for pathogenicity (Ma & Ma, 2016; McCann & Guttman, 2008). For example, *Xe* 85-10 harbors XopJ1/XopJ and XopJ3/AvrRxv that belong to the YopJ effector family and modify host cellular processes (Bonshtien et al., 2005; Üstun et al., 2013; Üstun & Börnke, 2015; Whalen et al., 2008). Other T3E families might be restricted, maybe emerging during adaptation to specific host. Most strains across the *Xanthomonas* genus harbor a set of 4 to 10 so-called core T3Es including AvrBs2, XopF1, XopK, XopL, XopM, XopN, XopP, XopQ, XopR, XopX and XopZ1 that are all present in the reference *Xe* 85-10 strain (Merda et al., 2017; Ryan et al., 2011; White et al., 2009). Several of these core T3Es impair plant defense responses and contribute to *Xanthomonas* spp. virulence in different hosts (Akimoto-Tomiyama et al., 2012; Li et al., 2015; Popov et al., 2016; Teper et al., 2016; Zhao et al., 2011). However, the functional characterization of individual T3E remains challenging due to complex interactions between T3Es and redundancy. The combined deletion of multiple T3Es or the generation of effectorless strains in *P. syringae* pv. DC3000 and *Ralstonia solanacearum* OE1-1 contributed to elucidate the diverse role of T3Es during bacterial infection (Lei et al., 2020; Wei et al., 2007; Wei et al., 2015). However this approach remain to be applied to *X. euvesicatoria* for a deeper understanding of virulence mechanisms.

The chemical control of leaf bacterial spot disease, relying on copper-based compounds and streptomycin, lost its efficiency with the widespread emergence of resistant *Xanthomonas* spp. strains (Stall et al., 2009). Crop protection strategies and breeding program shifted toward harnessing plant genetic resistance, through the identification, mapping and introgression of resistance loci. In pepper (*Capsicum* spp.), bacterial spot (*Bs*) resistance loci were identified from diverse genetic resources based on the hypersensitive response (HR), a form of programmed cell death triggered upon high titer inoculation of specific pathogen isolates (Jones et al., 2024; Stall et al., 2009). For example, *Capsicum chacoense Bs2* gene encodes an intracellular immune receptor belonging to the nucleotide-binding leucine-rich repeat receptor (NLR) family (Kourelis & van der Hoorn, 2018; Tai et al., 1999). Bs2 activation specifically occurs in response to *Xanthomonas* spp. strain carrying the conserved T3E avrBs2 and leads to HR and effector-triggered immunity (ETI) that effectively restricts bacterial growth (Ngou et al., 2022; Tai et al., 1999). Further, the NLRs Roq1 and NbZAR1 from *Nicotiana benthamiana* recognize the T3Es XopQ, XopJ1 and XopJ3, respectively and Roq1 transfer to tomato confers resistance to *Xe* 85-10 (Kim et al., 2026; Schultink et al., 2019; Thomas et al., 2020). *Bs2* was also successfully transferred to tomato, sweet orange (*Citrus sinensis*) and rice to confer resistance against *Xanthomonas* spp. harboring AvrBs2 (Du et al., 2025; Horvath et al., 2012; Kunwar et al., 2018; Sendín et al., 2017; Tai et al., 1999). However, polymorphic avrBs2 alleles can evade Bs2 recognition (Gassman et al., 2000; Horvath et al., 2012; Zhao et al., 2011). The breakdown of disease resistance provided by a single recognition event emphasizes the necessity of deploying multiple resistance genes/immune receptors for durable crop protection (Jayaraman et al., 2016; Oh & Choi, 2022; Shultink & Steinbrenner, 2021).

The black nightshade species *Solanum americanum* has recently emerged as a promising source of resistance. While cultivated *Solanum* species are susceptible to *P. syringae*, *R. solanacearum* or *Phytophthora infestans* infection, accessions of *S. americanum* exhibits a robust resistance, indicative of an expanded NLRs repertoire (Kim et al., 2025; Lin et al., 2023; Moon et al., 2021; Rodewald & Trognitz, 2013; Witek et al., 2016; Witek et al., 2021). Sequenced diploid homozygous *S. americanum* accessions are available for forward genetic screenings and amenable to Agrobacterium-mediated transformation, which eased the discovery of *P. syringae and R. solanacearum* effectors that trigger HR and of several *Resistance to P. infestans* (*Rpi*) genes (Kim et al., 2025; Lin et al., 2023; Moon et al., 2021; Witek et al., 2016; Witek et al., 2021). Here, we established *S. americanum* as a model system for investigating resistance to *X. euvesicatoria* infection. Using heterologous expression, multiple-effector knockout and complemented strains, we have identified nine *Xe* 85-10 T3Es (AvrBs2, XopAU, XopAP, XopE1, XopJ3, XopM, XopN, XopX and XopZ1) that trigger immune responses in *S. americanum* accession SP2273. More specifically, we could show that AvrBs2 and XopJ3 recognition by homologs of the corresponding NLRs Bs2 and ZAR1 was a major contributor to resistance against *X. euvesicatoria*. Furthermore, our approach revealed that AvrBs2 and XopJ3 were both required to enable bacterial multiplication when not recognized by the *S. americanum* immune system. This study brings insights into the important role of T3Es for bacterial multiplication in planta and provides a framework for the identification of immune receptors that may confer durable disease resistance in solanaceous crops.

## Results

### Effector-triggered immunity contributes to *S. americanum* resistance against *X. euvesicatoria*

To test *S. americanum* response to *X. euvesicatoria* infection, we first performed bacterial growth assays. *S. americanum* accession SP2273 was infected with a low inoculum of three *X. euvesicatoria* strains, the reference *Xe* 85-10 (Thieme et al., 2005), and two strains *Xe* 18722 (CNUPBL2030) and *Xe* 18723 (CNUPBL2058) isolated from pepper growth facilities in South Korea (Kyeon et al., 2016). At 3 and 6 days post inoculation (dpi), bacterial populations of the three strains remained at levels comparable to those at 0 dpi (Figure 1A), indicating that *X. euvesicatoria* cannot proliferate in *S. americanum*. The pathogenicity of the three strains was verified under laboratory conditions on a commercial tomato cultivar. Following dip-inoculation, characteristic leaf spot disease symptoms were observed in tomato plants (Supplementary Figure S1), confirming the pathogenic potential of the three strains.

**Figure 1.**
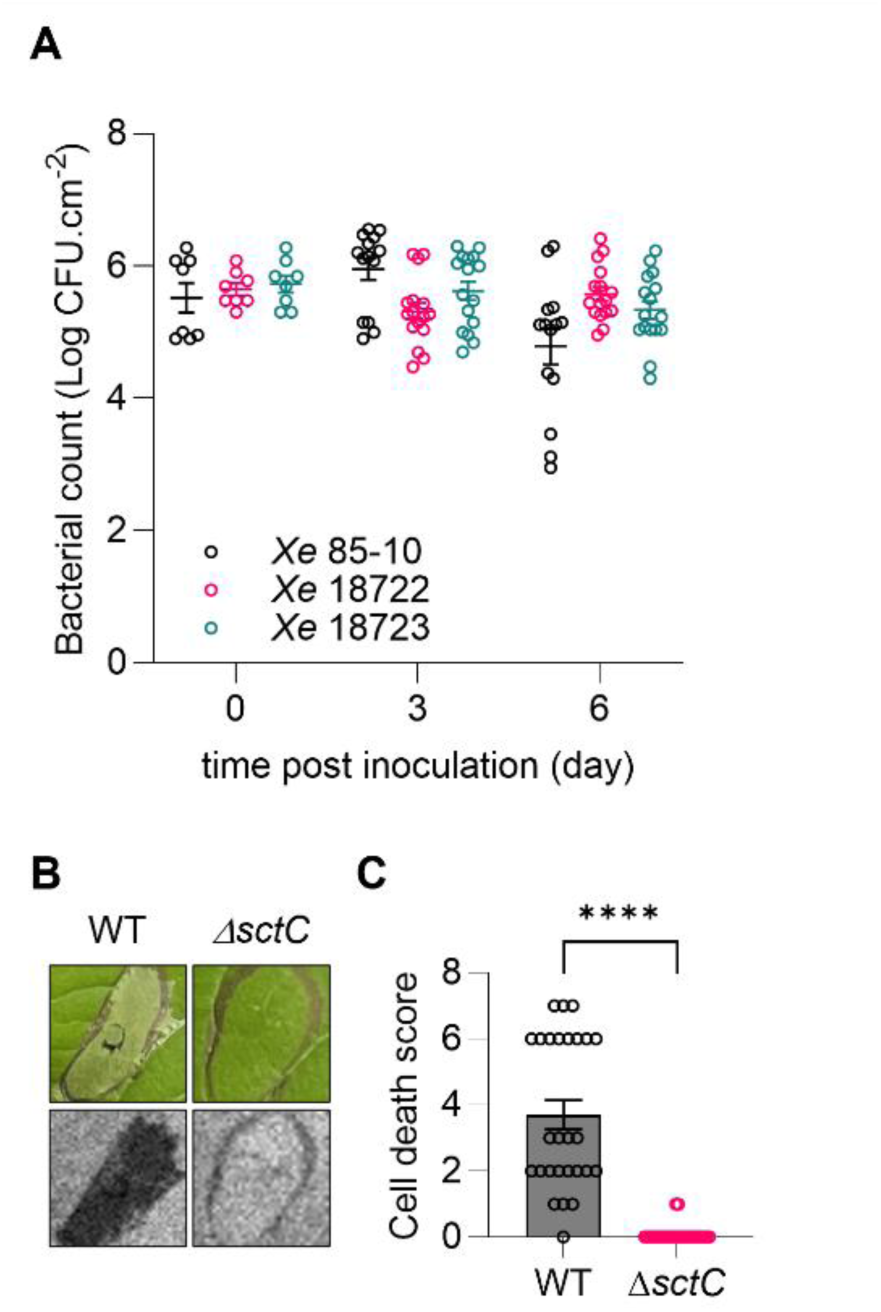
*X. euvesicatoria* strains do not grow and trigger hypersensitive response in *S. americanum* SP2273. A, Growth of *X. euvesicatoria* strains in *S. americanum* SP2273. Bacterial populations of *Xe* 85-10, *Xe* 18722 and *Xe* 18723 strains were quantified at the indicated time points after leaf infiltration with a low inoculum (OD 0.0005). Dots indicate individual values; bars represent mean +/- SEM from at least two biological repeats (n≥8). **B, C,** *X. euvesicatoria* triggers cell death in *S. americanum* SP2273. Leaves were infiltrated with a high inoculum (OD 0.05) of *Xe* 85-10 (WT) or *Xe* 85-10 type III secretion system knockout mutant (Δ*sctC*). Photographs (bright field on top panel, LED light on bottom panel) were taken 3 days after infiltration (**B**). Cell death intensity was evaluated on a programmed cell death range (**C**). Dots indicate individual values; bar graphs represent mean +/- SEM from five biological repeats (n=35). Asterisks indicate statistical difference between WT and Δ*sctC* strains (Student’s t- test; ****, *P*<0.0001).

We next investigated whether the restriction of *X. euvesicatoria* growth in *S. americanum* is mediated by effector-triggered immunity (ETI). The *Xe* 85-10 strain and the T3SS-deficient mutant *Xe* 85-10 Δ*sctC* (Portaliou et al., 2016) were infiltrated at high titer in *S. americanum* SP2273 leaf. In contrast to *Xe* 85-10 Δ*sctC*, *Xe* 85-10 induced a rapid and robust hypersensitive response (HR)-like cell death by 2 dpi (Figure 1B). Quantitative assessment of the cell death using the programmed cell death range described by Kim et al. (2025) further confirmed the significant difference in the intensity of the response (Figure 1C). Taken together, these results suggest that the T3Es of *Xe* 85-10 trigger ETI and contribute to *S. americanum* resistance against *X. euvesicatoria* infection.

### AvrBs2 and XopJ3 are major contributors to ETI in *S. americanum*

To identify the *Xe* 85-10 effectors that activate ETI in *S. americanum*, 37 of the 38 known T3Es were transiently expressed in SP2273 leaf (Schulze et al., 2012; Thieme et al., 2005; Teper et al., 2016). Ten effectors, AvrBs2, XopAP, XopAU, XopE1, XopF2, XopJ3, XopM, XopN, XopX, and XopZ1, triggered typical HR-like cell death by 5 dpi (Supplementary Figure S2A). Quantitative assessment of the cell death further confirmed a significant difference in the response between each effectors and the negative control GFP (Supplementary Figure S2B), indicating that these ten effectors might be recognized by the *S. americanum* immune system. The accumulation of the remaining 27 effectors that did not trigger cell death was confirmed by immunoblotting (Supplementary Figure S3).

Since effector expression from a constitutive promoter differs from the delivery by the T3SS, we sequentially knocked out nine recognized effectors from *Xe* 85-10 to evaluate their contribution in a more natural context. As avrBs2 and XopJ3 are recognized in solanaceous species by Bs2 and NbZAR1, respectively (Kim et al., 2026; Tai et al., 1999), we first generated the Δ*avrBs2* Δ*xopJ3* (D2E) strain using an homologous recombination system (Supplementary Figure S4). The remaining recognized effectors were then knocked out sequentially to generate D6E (D2E Δ*xopAU* Δ*xopM* Δ*xopN* Δ*xopZ1*) and D9E (D6E Δ*xopAP* Δ*xopE1* Δ*xopX*). To assess the ETI-eliciting ability of multiple effector knock-out mutants, *Xe* 85-10 wild-type (WT), Δ*sctC*, D2E, D6E, and D9E strains were infiltrated at high titer in SP2273 leaf. Compared to *Xe* 85-10 WT, D2E induced moderately reduced cell death, D6E showed a further reduction, and D9E failed to induce any detectable cell death, mirroring the phenotype of *Xe* 85-10 Δ*sctC* (Figure 2). Additionally, we quantified the expression of the ETI-marker gene *SaFMO1* in the same conditions (Mishina & Zeier, 2006). Consistent with the cell death assays, *SaFMO1* expression was significantly lower in leaf inoculated with D6E and D9E compared to leaf infiltrated with *Xe* 85-10 WT (Supplementary Figure S5). These results suggest that at least 9 T3Es collectively contribute to ETI activation in *S. americanum*.

**Figure 2.**
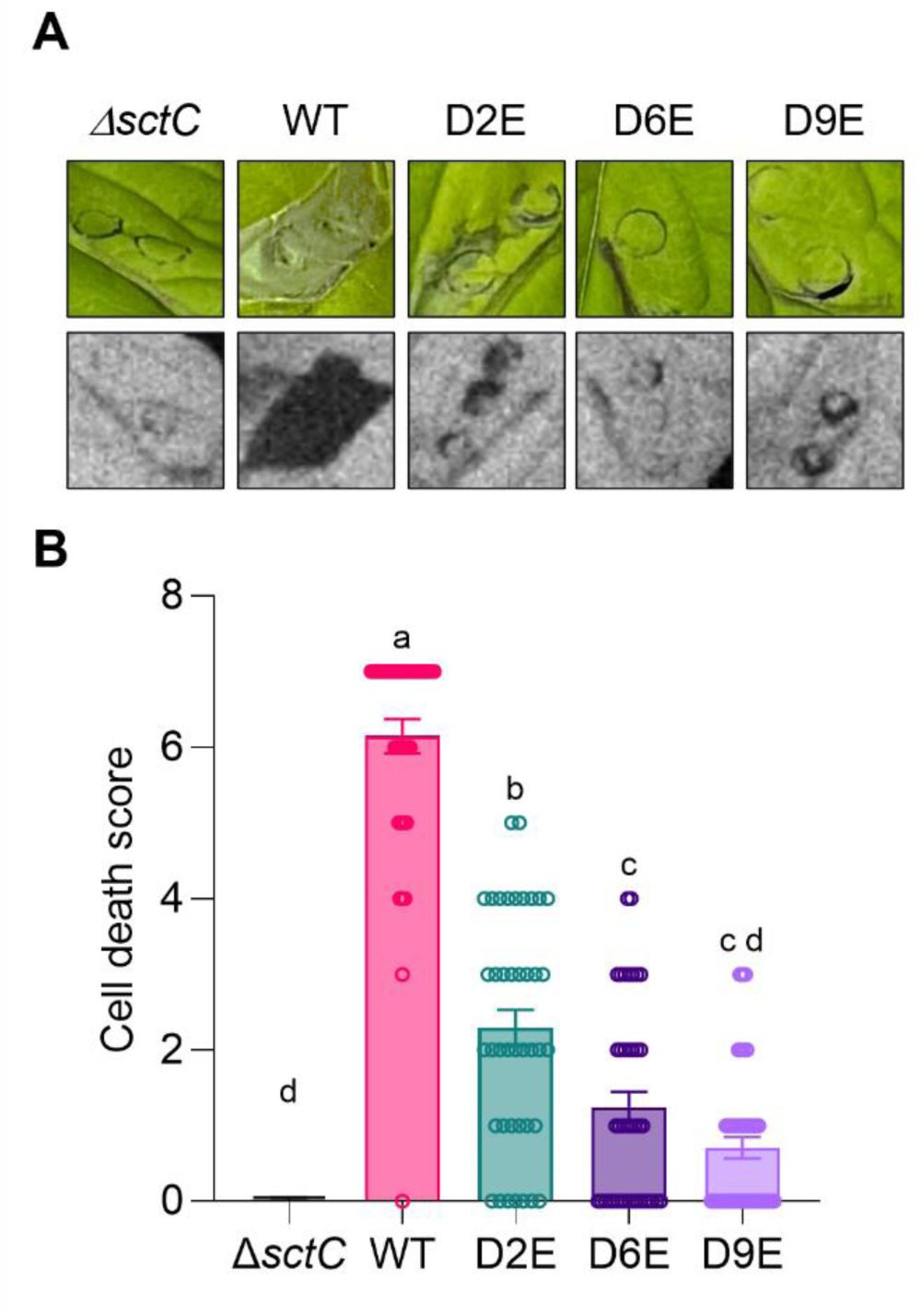
Knockout of nine effectors abolishes the cell death triggered by *Xe* 85-10 in *S. americanum* SP2273. A, B, Cell death triggered by *Xe* 85-10 WT, Δ*sctC*, D2E (Δ*avrBs2* Δ*xopJ3*), D6E (Δ*avrBs2* Δ*xopJ3* Δ*xopAU* Δ*xopM* Δ*xopN* Δ*xopZ1*), and D9E (Δ*avrBs2* Δ*xopJ3* Δ*xopAU* Δ*xopM* Δ*xopN* Δ*xopZ1*Δ *xopAP* Δ*xopE1* Δ*xopX*) in *S. americanum* SP2273. Photographs (bright field on top panel, LED light on bottom panel) were taken 2 to 4 days after infiltration (**A**). Cell death intensity was evaluated on a programmed cell death range (**B**). Dots indicate individual values; bar graphs represent mean +/- SEM from five biological repeats (n=35). Different letters indicate statistical difference between strains (one-way ANOVA, Tukey’s post hoc test; *P*<0.05).

To determine the individual contribution of the nine T3Es when delivered via the native T3SS, each effector gene was reintroduced in the D9E strain on a broad host range plasmid (Kovach et al., 1995). The nine D9E complementation strains along with D9E (EV) and *Xe* 85-10 WT (EV) were infiltrated in SP2273 at high titer. When delivered by D9E, AvrBs2 and XopJ3 restored a robust HR-like cell death, while XopAP and XopN induced a moderate cell death response (Figure 3). However, only the D9E delivery of AvrBs2 and to a lesser extent of XopJ3 could increase *SaFMO1* expression (Supplementary Figure S6). Individual effector expression in the D9E complemented strains was confirmed by immunoblotting (Supplementary Figure S7). These findings demonstrate that while nine effectors may contribute collectively and/or partially to the recognition of *X. euvesicatoria*, AvrBs2 and XopJ3 are major determinants of ETI in *S. americanum*.

**Figure 3.**
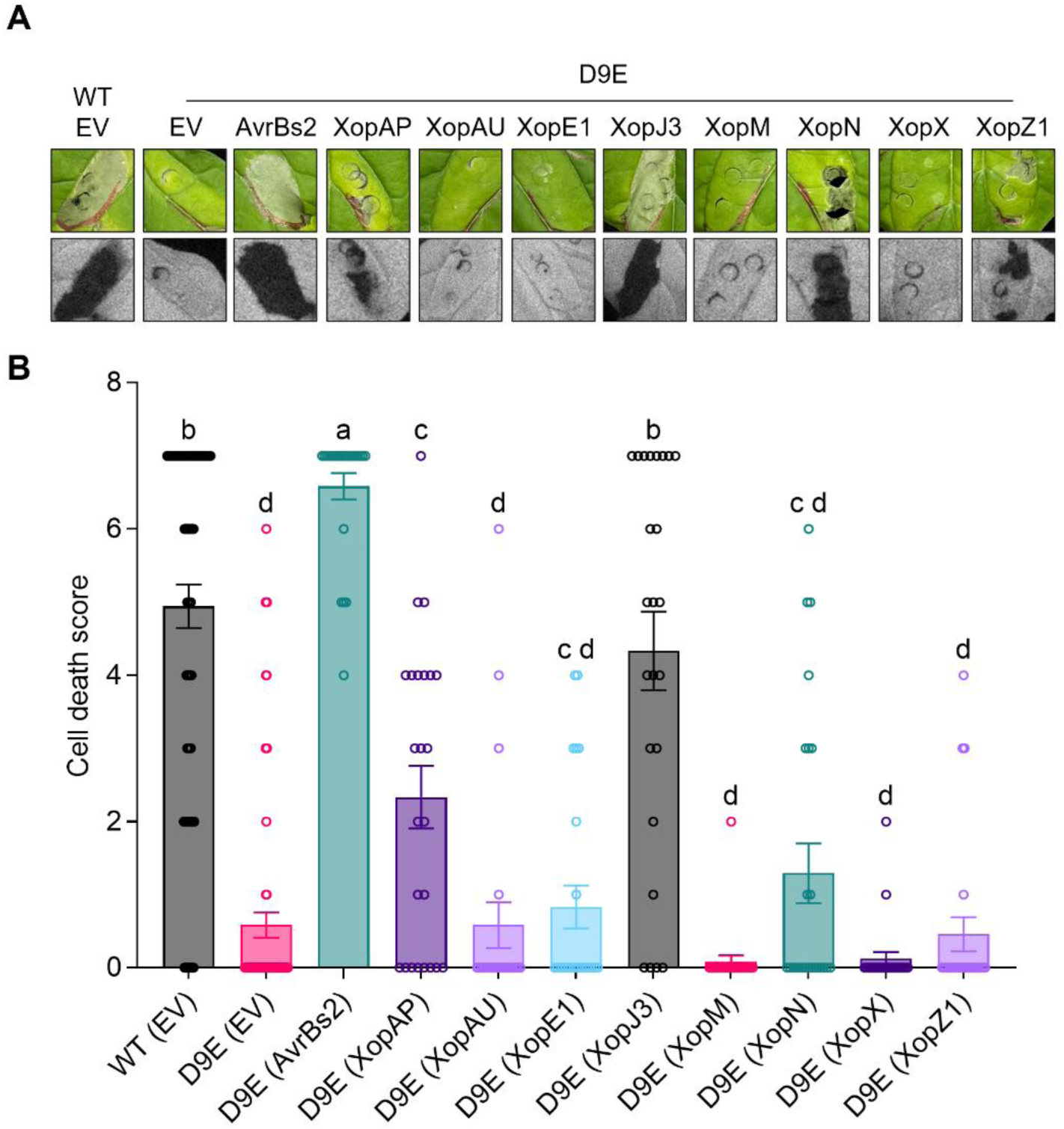
Complementation of *Xe* DGE strain with AvrBs2 or XopJ3 restores cell death in *S. americanum*. A, B, Cell death triggered by *Xe* 85-10 WT, D9E and D9E complemented strains (D9E(EV), D9E(AvrBs2), D9E(XopAP), D9E(XopAU), D9E(XopE1), D9E(XopJ3), D9E(XopM), D9E(XopN), D9E(XopX), and D9E(XopZ1)) in *S. americanum* SP2273. Photographs (bright field on top panel, LED light on bottom panel) were taken 3 days after infiltration (**A**). Cell death intensity was evaluated on a programmed cell death range (**B**). Dots indicate individual values; bar graphs represent mean +/- SEM from five biological repeats (n=30). Different letters indicate statistical difference between strains (one-way ANOVA, Tukey’s post hoc test; *P*<0.05).

### *SaBs2* and *SaZAR1* homologs recognize AvrBs2 and XopJ3, respectively

Given that AvrBs2 and XopJ3 were major ETI elicitors in *S. americanum*, we investigated the presence and function of *Bs2* and *NbZAR1* homologs in the SP2273 genome (Lin et al., 2023). A BLASTn search using the *Capsicum chacoense Bs2* gene and *N. benthamiana ZAR1* sequences as queries identified four *Bs2* homologs, denoted *SaBs2a* (SP2273_Chr11_NLR96), *SaBs2b* (SP2273_Chr12_NLR01), *SaBs2c* (SP2273_Chr12_NLR40), and *SaBs2d* (SP2273_Chr12_NLR41) and one *NbZAR1* homolog, *SaZAR1* (SP2273_Chr02_NLR08), based on previously established nomenclature (Tai et al., 1999; Ahn et al., 2023; Lin et al., 2023). The predicted SaZAR1 protein exhibits high amino acid sequence identity with NbZAR1 (89 %; Supplementary Figure S8A). Protein sequence alignments also revealed that the four *SaBs2* homologs share high amino acid identity with Bs2 (55-67%; Supplementary Figure S8A). Furthermore, expression analysis indicated that *SaZAR1* and *SaBs2* homolog transcripts, with the exception of *SaBs2c*, could accumulate in SP2273 leaf (Figure S8B). Therefore, we generated a quintuple *SaZAR1*-*SaBs2a*-*SaBs2b*-*SaBs2c*-*SaBs2d* knockout line using CRISPR/Cas9-based multiplex gene editing system (Stuttman et al., 2021). Edition sites were targeted by 16 sgRNAs (4 for *SaZAR1* and 3 for each *SaBs2* gene), and successful knockout was confirmed by Sanger sequencing (Supplementary Figure S9A). Among the 73 primary T_0_ transformants regenerated via tissue culture, four lines (T_0_#15, #32, #39, #42) exhibited impaired cell death in response to AvrBs2 and HopZ1a+ZED1 (Lewis et al., 2013) (Supplementary Figure S9B). The specific loss of cell death in these four T_0_ lines, which carried homozygous or heterozygous mutations in the five target genes, suggested that the editing of *SaZAR1* and *SaBs2* homologs, rather than off-target mutations, directly impaired the effector recognition. Line #42 was advanced and fixed at the T_2_ generation as a homozygous quintuple mutant (hereafter referred to as the SP2273-*bz* line).

Next, we transiently expressed AvrBs2 and XopJ3 in SP2273 and SP2273-*bz* via Agrobacterium-mediated transformation. GFP and Avramr1 (recognized by *Rpi-amr1*; Witek et al., 2021) served as negative and positive controls, respectively. While AvrBs2 and XopJ3 triggered robust HR-like cell death in SP2273, this response was completely abolished in 2273-*bz* (Figure 4A and B). Importantly, 2273-*bz* maintained the cell death response to Avramr1, confirming that the loss of recognition for AvrBs2 and XopJ3 was specific to the knockout of *SaBs2* and *SaZAR1* rather than a general defect in immune signaling. To further validate these results, we performed genetic complementation assays. Co-expression of XopJ3 and SaZAR1 restored the cell death response in SP2273-*bz* (Figure 4C and D). Co-expression of AvrBs2 with each of the four SaBs2 homologs successfully restored HR-like cell death (Figure 4E and F), indicating that the SaBs2 homologs display redundant AvrBs2 recognition ability, at least for *SaBs2a*, *SaBs2b* and *SaBs2d*, which are expressed in SP2273 leaf.

**Figure 4.**
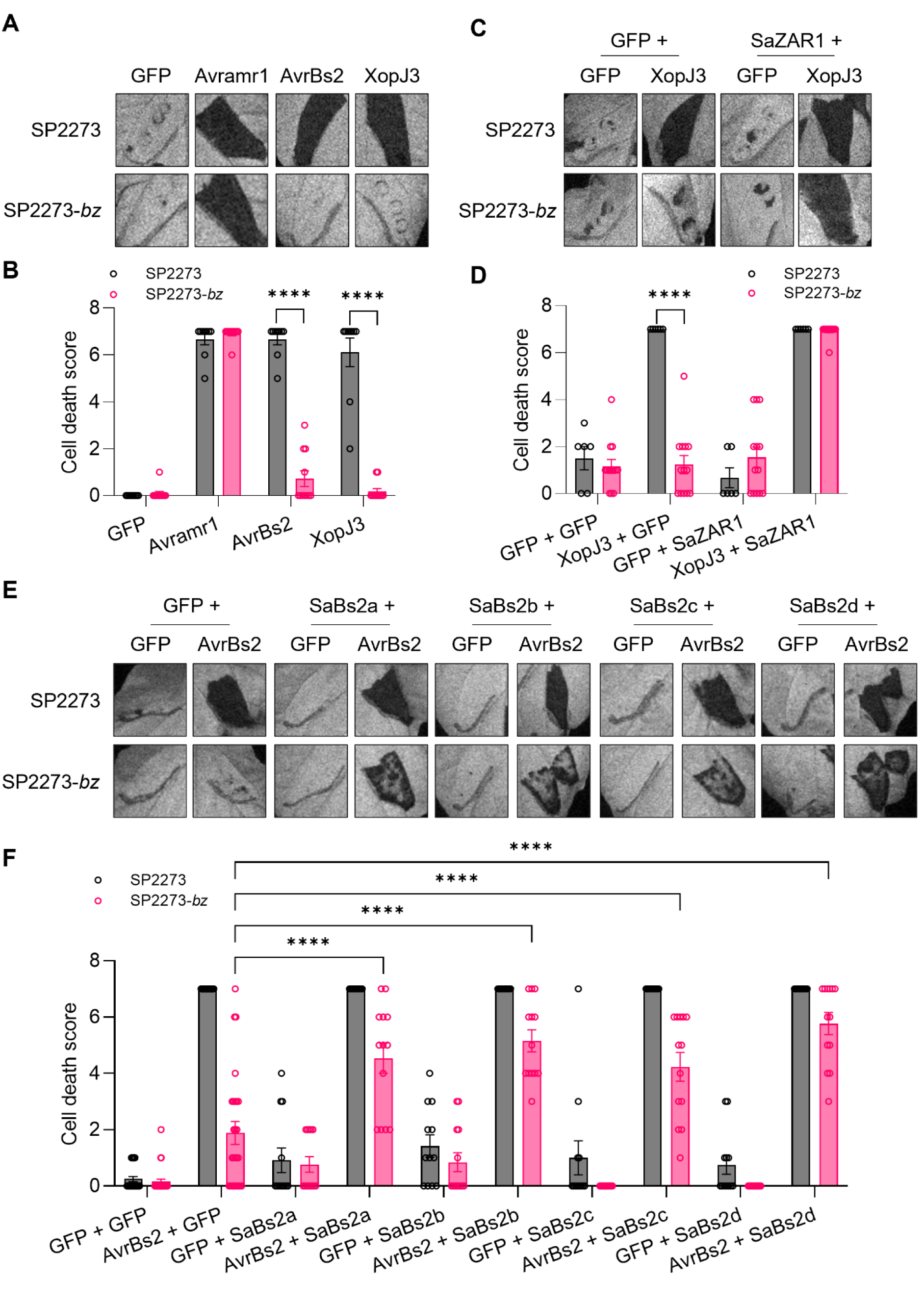
AvrBs2 and XopJ3 are recognized by *S. americanum* homologs of *Bs2* and *ZAR1*, respectively. A, B, Cell death triggered by Agrobacterium-mediated expression of GFP, AvrBs2, XopJ3 or Avramr1 in *S. americanum* SP2273 or SP2273 line edited at *SaBs2a*, *SaBs2b*, *SaBs2c*, *SaBs2d* and *SaZAR1* loci (SP2273-*bz*). **C, D,** Cell death triggered by Agrobacterium-mediated co-expression of GFP or XopJ3 with GFP or SaZAR1 in *S. americanum* SP2273 or SP2273-*bz*. **E, F,** Cell death triggered by Agrobacterium-mediated co-expression of GFP or AvrBs2 with GFP, SaBs2a, SaBs2b, SaBs2c or SaBs2d in *S. americanum* SP2273 or SP2273-*bz*. Photographs (LED light) were taken 3 days after infiltration (**A, C, E**). Cell death intensity was evaluated on a programmed cell death range (**B, D, F**). Dots indicate individual values; bar graphs represent mean +/- SEM from at least two biological repeats (n≥9 in **B**; n≥6 in **D**; n≥12 in **F**). Asterisks indicate statistical differences between the two plant genotypes in **B** and **D** or with the AvrBs2+GFP condition in the SP2273-*bz* line in **F** (two-way ANOVA, Šidák’s multiple comparisons; ****, *P*<0.0001).

We also tested the contribution of AvrBs2 and XopJ3 recognition to the *S. americanum* immune response to *X. euvesicatoria* infection. We inoculated SP2273 and SP2273-*bz* plants with *Xe* 85-10 WT, Δ*avrBs2*, Δ*xopJ3*, D2E, D9E and Δ*sctC* strains at a high titer. While *Xe* 85-10 WT induced robust HR-like cell death in SP2273, this response was significantly reduced in SP2273-*bz* (Figure 5). In accordance with previous results, D2E and D9E strains did not elicit cell death in SP2273 and SP2273-*bz*. Interestingly, SP2273 response to Δ*avrBs2* but not to Δ*xopJ3* was reduced compared to the response to *Xe* 85-10 WT, suggesting that AvrBs2 recognition has a major role in eliciting cell death in *S. americanum*. However, SP2273-*bz* responses to Δ*avrBs2* and Δ*xopJ3* were significantly reduced compared to SP2273, indicating that *SaBs2* and *SaZAR1* homologs collectively contribute to the cell death elicited by *X. euvesicatoria* in *S. americanum*. *X. vesicatoria* (formerly *X. campestris* pv. *vesicatoria*) *Xcv* Bv5-4, carrying the independently recognized effector XopJ2/AvrBsT (Kim et al., 2010; Sharma et al., 2024), was used as an additional control. *Xcv* Bv5-4 induced cell death in SP2273 and in SP2273-*bz*, confirming that SP2273-*bz* remains capable of AvrBsT recognition (Figure 5). Taken together, these results demonstrate that the recognition of AvrBs2 by *SaBs2* and XopJ3 by *SaZAR1* is the major component of ETI in S*. americanum*.

**Figure 5.**
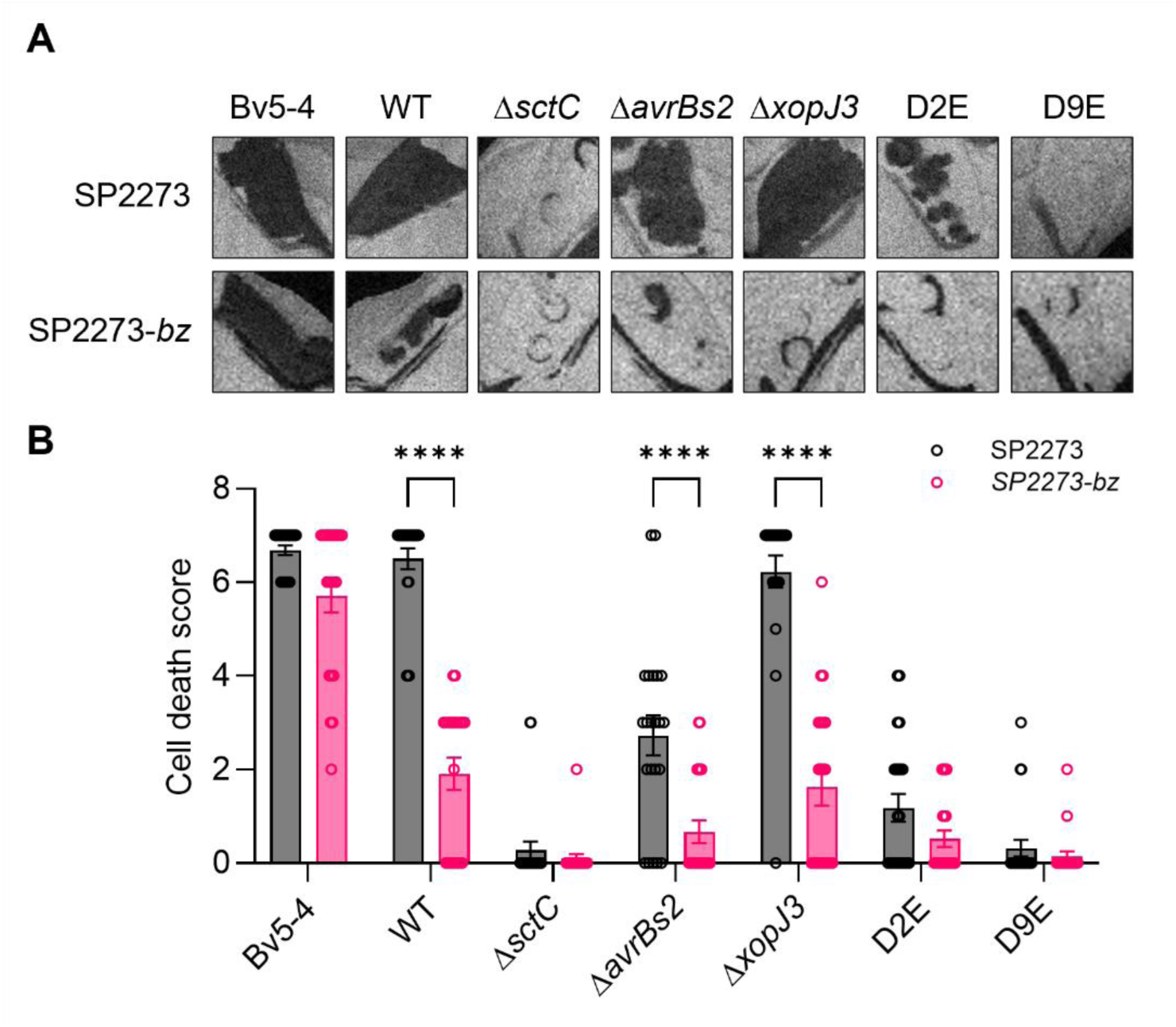
*X. euvesicatoria*-triggered cell death is impaired in SP2273-*bz* line. A, B, Cell death triggered by *Xcv* Bv5-4, *Xe* 85-10 WT, Δ*sctC*, Δ*avrBs2*, Δ*xopJ3*, Δ*avrBs2* Δ*xopJ3* (D2E) or D9E strains in *S. americanum* SP2273 or SP2273-*bz*. Photographs (LED light) were taken 3 days after infiltration (**A**). Cell death intensity was evaluated on a programmed cell death range (**B**). Dots indicate individual values; bar graphs represent mean +/- SEM from three biological repeats (n=22). Asterisks indicate statistical differences between the two plant genotypes (two-way ANOVA, Šidák’s multiple comparisons; ****, *P*<0.0001).

AvrBs2 and XopJ3 are both required for *X. euvesicatoria* infection in *S. americanum*

To evaluate how the loss of AvrBs2 and XopJ3 recognition affects bacterial multiplication *in planta*, we compared bacterial growth in SP2273 and SP2273-*bz*. At 6 dpi, *Xe* 85-10 WT population in SP2273- *bz* was significantly higher than in SP2273 (Figure 6A), suggesting that AvrBs2 and XopJ3 recognition restricted bacterial growth in *S. americanum*. Interestingly, Δ*avrBs2*, Δ*xopJ3*, and D2E strains exhibited restricted growth compared to *Xe* 85-10 WT in SP2273-*bz*, indicating that AvrBs2 and xopJ3 recognition collectively limit *X. euvesicatoria* infection. These results also suggest that AvrBs2 and XopJ3 possess virulence functions that promote *X. euvesicatoria* multiplication in *S. americanum*.

**Figure 6.**
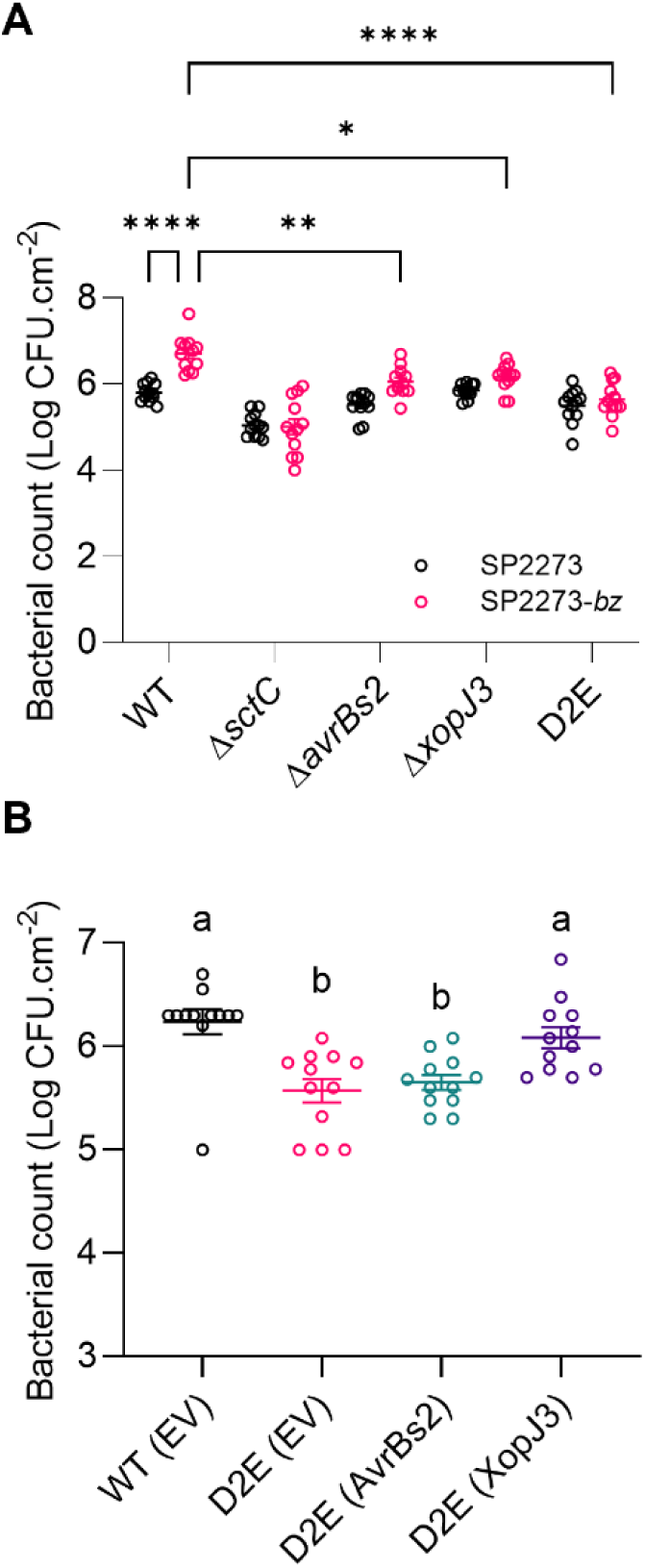
AvrBs2 and XopJ3 both contribute to *X. euvesicatoria* growth in *S. americanum* SP2273. A, Growth of *Xe* 85-10 WT, Δ*sctC*, Δ*avrBs2*, Δ*xopJ3*, and Δ*avrBs2* Δ*xopJ3* (D2E) strains in *S. americanum* SP2273 or SP2273-*bz*. **B,** Growth of *Xe* 85-10 WT, D2E and D2E complemented with AvrBs2 or XopJ3 in *S. americanum* SP2273-*bz*. Bacterial populations were quantified 6 days after leaf infiltration with a low inoculum (OD 0.0005). Dots indicate individual values; bars represent mean +/- SEM from three biological repeats (n=12). Asterisks indicate statistical differences with the *X. euvesicatoria* 85-10 WT in SP2273-*bz* condition in **A** (two-way ANOVA, Šidák’s multiple comparisons; *, *P*<0.05; **, *P*<0.01; ****, *P*<0.0001). Different letters indicate statistical differences between strains in **B** (one-way ANOVA, Tukey’s post-hoc test; *P*<0.05).

To assess AvrBs2 and XopJ3 contribution to *X. euvesicatoria* virulence, each effector was reintroduced in the D2E strain for growth assay in SP2273-*bz*. While D2E (EV) and D2E (AvrBs2) strains showed significantly decreased growth relative to *Xe* 85-10 WT (EV) in SP2273-*bz*, D2E (XopJ3) populations were similar to *Xe* 85-10 WT (Figure 6B). These results suggest that while AvrBs2 and XopJ3 both contribute to *X. euvesicatoria* infection, XopJ3 may play a more significant role in promoting bacterial growth. Altogether, our findings highlight a dual role for AvrBs2 and XopJ3, as factors essential for *X. euvesicatoria* multiplication and, simultaneously, as major determinants of ETI in *S. americanum*.

## Discussion

Our work identified multiple *Xe* 85-10 effectors that elicit immune responses in the model solanaceous species *S. americanum*. Generation of multiple effector knockout strains and quantification of immune responses demonstrated that nine *Xe* 85-10 effectors trigger ETI in *S. americanum* and suggest that ETI plays a major role in *S. americanum* resistance to *X. euvesicatoria* infection. Recognition of multiple effectors is a mechanism underlying nonhost resistance as demonstrated for *Phytophthora infestans* in pepper or *Pseudomonas syringae* in *S. americanum* for example (Kim et al., 2025; Lee et al., 2014). We acknowledge that our experimental approach bypassed pre-invasive barriers that contribute to nonhost resistance. While syringe infiltration effectively isolates post-invasive defense mechanisms, it may not fully capture the complexity of nonhost resistance. Beside multiple effector recognition events, the role of *S. americanum* cell wall composition for bacterial attachment and the presence of surface-localized immune receptors and/or inhibitory metabolites should additionally be considered (Lee et al., 2017; Panstruga & Moscou, 2020; Wu et al., 2023). Nevertheless, the distinct immune responses elicited by *Xe* 85-10 WT and T3SS or multiple T3E mutants, as well as the enhanced proliferation of *Xe* 85-10 in the SP2273-*bz* line provide robust evidence that ETI is the primary contributor to *S. americanum* resistance.

Moreover, the generation of *S. americanum* plants lacking SaBs2 and SaZAR1 receptors revealed the ambivalent role of AvrBs2 and XopJ3. As indicated by the reduced growth of D2E compared to the *Xe* 85-10 WT in SP2273-*bz*, both effectors appear essential for *X. euvesicatoria* infection in *S. americanum*. AvrBs2 is highly conserved in Xanthomonads and is required for full virulence in susceptible hosts such as alfafa and pepper (Kearney & Staskawicz, 1990). A recent study further revealed the catalytic function of AvrBs2 in xanthosan synthesis in *X. oryzae*, *X. citri* and *X. euvesicatoria* (Wang et al., 2026). In regards, the inability of the Δ*avrbs2* strain to grow to the same levels as *Xe* 85-10 WT in SP2273-*bz* observed in this study could likely stem from an impairment in nutrient uptake and further supports the interest of developing “anti-nutrition” strategies for bacterial disease control in crops (Mahmood et al., 2026; Phan et al., 2026; Wang et al., 2026).

However, we also observed that the complementation of the D2E strain with avrBs2 alone could not restore higher bacterial populations in SP2273-*bz*, while D2E (xopJ3) achieved higher growth. XopJ3 belongs to the conserved YopJ family of bacterial effectors comprising the well-characterized acetyltransferase *R. solanacearum* RipP2/PopP2 and *P. syringae* HopZ5 (Ma & Ma, 2016). These T3Es contribute to bacterial virulence through the inhibitory acetylation of specific components of the plant immune system such as WRKY transcription factors and ARID subunits of chromatin remodeling complexes for RipP2, or the NOI-containing disordered protein RIN4 (Choi et al., 2021; Le Roux et al., 2015; Monge-Waleryszak, 2025). XopJ3/AvrRxv was first characterized as an avirulence factor in bean and tomato (Whalen et al., 1993). XopJ3 also triggers NbZAR1-dependent ETI in *N. benthamiana* (Albers et al., 2019; Kim et al., 2026). While the conserved predicted catalytic Cys244 is essential for XopJ3 recognition by NbZAR1, the function and possible substrates of XopJ3 in host cells remain to be elucidated (Kim et al., 2026; Whalen et al., 2008). It is possible that XopJ3, like RipP2 or HopZ5, has an immunosuppressive role, which could facilitate the delivery or hinder the recognition of other T3Es – including the seven other T3E identified here. Together, the delivery of both AvrBs2 and XopJ3 in cells lacking the corresponding immune receptors would allow *X. euvesicatoria* growth in *S. americanum* through combined immunosuppression and nutrient acquisition activities. The successful multiplex knockout of 5 NLRs in *S. americanum* genome also supported the potential of this species as an interesting model for studies on NLR evolution. In *N. benthamiana*, NbZAR1 and the accessory receptor-like cytoplasmic kinase (RLCK) JIM2 recognize multiple YopJ family effectors including Xanthomonas spp. XopJ1/XopJ, XopJ2/AvrBsT, XopJ3/AvrRxv and XopJ4 (Ahn et al., 2023; Kim et al., 2026; Schultink et al., 2019). *Xe* 85-10 harbors two members of the YopJ family, XopJ1 and XopJ3 (Schulze et al., 2012; Thieme et al., 2005; Teper et al., 2016) but our assay revealed that only XopJ3 could trigger ZAR1-dependent immune responses in *S. americanum*. Hence, despite the high homology with NbZAR1, SaZAR1 appeared to display a distinctive spectrum of recognition. Genomic analyses of the RLCK family in *S. americanum* would provide further support to characterize SaZAR1 mechanism of activation and possibly strengthen evidence for the co-evolution of this ancient immune receptor with the associated regulatory kinases (Adachi et al., 2023; Diplock et al., 2022; Gong et al., 2022).

The genomic architecture of *S. americanum* can be distinguished from that of cultivated relatives by the presence of multiple *Bs2* homologs. While pepper (*Capsicum chacoense*) and tomato (*S. lycopersicum*) contain a single *Bs2* gene (not functional in tomato; Tai et al., 1999), we identified 4 loci with high homology to *Bs2* in *S. americanum* SP2273 genome (Bolger et al., 2014; Lin et al., 2023). Sequence and structure prediction (not shown) analyses confirmed the canonical functional CNL structure for the proteins coded by these 4 loci, while transient expression assays in SP2273-*bz* demonstrated that each SaBs2 homolog functions for AvrBs2 recognition. Although we could not detect SaBs2c transcript in SP2273 leaf tissues, our results indicate that the 4 *SaBs2* genes are functional and possibly redundant. The presence of multiple functional *Bs2* copies in *S. americanum* may participate in a broad-spectrum surveillance strategy. Considering that Xanthomonas spp. field isolates harbor polymorphic avrBs2 that can escape recognition while conserving the virulence function (Gassman et al., 2000; Horvath et al., 2012; Zhao et al., 2011), the maintenance of 4 *Bs2* homologs in *S. americanum* genome may counteract the rapid effector evolution. Further characterization of functional polymorphisms among *SaBs2* genes may inform the status and editing potential of nonfunctional *Bs2* homologs, which remain poorly understood. Additionally, the functional disparity of AvrBs2 recognition between pepper, tomato and *S. americanum* may be further determined by differences in the helper NRC network required downstream of *Bs2* activation (Liu et al., 2024; Wu et al., 2017). Elucidating the molecular basis for the loss of function in Bs2 and NRC homologs could enable precision genome editing and engineering of durable resistance in susceptible crops hopefully with similar success as the combined transfer of *Bs2* and *NbNRCs* in rice (Du et al., 2025).

Lastly, the identification of T3Es recognized by a resistant model species constitute a valuable tool for the discovery of immune receptors and effector-assisted breeding (Jayaraman et al., 2016; Vleeshouwers & Oliver 2014). While the receptors for AvrBs2 and XopJ3 were identified in this work through genomic analyses, the NLRs involved in the recognition of the remaining T3Es (XopAU, XopAP, XopE1, XopM, XopN, XopX and XopZ1) can be mapped using forward genetics, leveraging the natural variation within *S. americanum*. Together, these approaches could enable the engineering of durable resistance in susceptible crops through the precise manipulation of core effector recognition.

## Materials and Methods

### Molecular constructs

The coding sequences of 37 effectors retrieved from the reference strain *X. euvesicatoria* 85-10 (Seong et al., 2016; Thieme et al., 2005) were divided into 1 to 1.5 kb modules. Module DNA was amplified from *Xe* 85-10 genomic DNA with the flanking BsaI site-containing primers listed in Supplementary Table S1. PCR products were ligated into the entry vector pICH41021 and each module construct was verified by Sanger sequencing. Entry modules were assembled into the binary vector pICH86988 in fusion with a C-terminal 3xFLAG tag under the Cauliflower Mosaic Virus 35S promoter using the Golden Gate cloning method (Engler et al., 2008). Assemblies confirmed by restriction analysis were mobilized into *Agrobacterium tumefaciens* AGL1 strain by electroporation. Entry modules for avrBs2, xopAP, xopAU, xopE1, xopJ3, xopM, xopN, xopX and XopZ1 were also assembled into the broad host-range vector pBBR1 in fusion with a C-terminal 3xFLAG tag under the control of the xopO promoter (Kovach et al., 1995; Koebnik et al., 2006) and mobilized into the D9E strain by triparental mating using *Escherichia coli* HB101 helper strain, as described previously (Jayaraman et al., 2017).

For bacterial gene deletion, upstream and downstream flanking regions (∼1000 bp) of sctC, avrBs2, xopAP, xopAU, xopE1, xopJ3, xopM, xopN, xopX and XopZ1 were amplified from *Xe* 85-10 genomic DNA with primers listed in Supplementary Table S1 and ligated into the entry vector pICH41021. Sequenced modules were assembled into the pK18mobsacB vector modified for Golden Gate (Jayaraman et al., 2020) and mobilized into *E. coli* DH5α strain by electroporation.

For genetic complementation assays, *SaBs2a*, *SaBs2b*, *SaBs2c*, *SaBs2d* and *SaZAR1* coding sequences retrieved from *S. americanum* SP2273 genome (Lin et al., 2023) were amplified as ∼ 1 kb modules with primers listed in Supplementary Table S1 and cloned into pICH41021. Sequenced modules were assembled into the binary vector pICH86988 in fusion with a C-terminal HA or 6xHA (for *SaZAR1*) tag under the Cauliflower Mosaic Virus 35S promoter. Assemblies were mobilized into *Rhizobium rhizogenes* AS109 (Lopez-Agudelo et al., 2025) or *A. tumefaciens* AGL1 (for *SaZAR1*) strain by electroporation.

### Plant growth conditions

*S. americanum* accession SP2273 was used in this study (Lin et al., 2023). Seeds were incubated in 2.5 mM gibberellic acid for 24 h at 28°C and sowed on mixed commercial soil (Seoul Bio, Republic of Korea). *S. americanum* plants were grown in a chamber at 25°C in short-day conditions (13 h light/11 h dark) for 5 to 6 weeks before experiments. *N. benthamiana* and commercial tomato cultivar Lovely 256 (Asia Seeds, Republic of Korea) were grown in a chamber at 25°C in short-day conditions for 4 to 5 weeks before experiments.

### Bacteria growth conditions

*Xe* 85-10 (Seong et al., 2016) and *Xcv* Bv5-4 (Kim et al., 2010) were kind gifts from Prof. Sang-Wook Han (Chung-Ang University, Anseong, Republic of Korea). *Xe* 18722 and *Xe*18723 (Kyeon et al., 2016) were obtained from the Korean Agricultural Culture Collection. *X. euvesicatoria* strains were grown in NYG (peptone yeast glycerol) medium containing appropriate antibiotics (rifampicin at 20 mg/l for *Xe* 85-10; gentamycin at 20 mg/l for *Xe* 18722 and *Xe* 18723) for 48 to 72 h at 28°C. Single colonies were transferred to liquid Luria-Bertani broth (LB) containing the same antibiotics and cultured for 18 to 24 h at 28°C before experiments. *Agrobacterium tumefaciens* AGL1 strains were grown on solid LB medium containing carbenicillin at 100 mg/ml and kanamycin at 50 mg/l for 24 to 48 h at 28°C. Single colonies were transferred to liquid LB medium and cultured for 18 to 24 h at 28°C before experiments. *R. rhizogenes* AS109 strains were grown on solid 523 medium containing rifampicin at 20 mg/l and kanamycin at 50 mg/l for 72 to 96 h at 28°C. Single colonies were transferred to liquid 523 medium containing the same antibiotics and cultured for 36 to 48 h at 28°C before experiments.

### Bacterial leaf spot disease assay in tomato

*X. euvesicatoria* cells were harvested by centrifugation at 3,000 *g* for 3 minutes and resuspended in 10 mM MgCl_2_. Tomato plants were dip-inoculated for 15 seconds in 10 mM MgCl_2_ or bacterial suspensions (OD_600_ 0.00001) (Gu et al., 2011). Tomato plants were covered with transparent plastic bag for 24 h and kept at room temperature (23-25°C) for 15 days.

### Cell death assay

*X. euvesicatoria* (in 10 mM MgCl_2_; OD_600_ 0.05), *A. tumefaciens* (in 10 mM MgCl_2_, 10 mM MES-KOH pH 5.6; OD_600_ 0.4) or *R. rhizogenes* (in 10 mM MgCl2, 10 mM MES-KOH pH 5.6, 200 μM acetosyringone; OD_600_ 0.5) strains were infiltrated into *S. americanum* leaf using needleless syringes. Cell death intensity was quantified manually using the programmed cell death range reported by Kim et al. (2025) or by measuring the quantum yield of chlorophyll fluorescence (Fv/Fm) using a closed FluorCam (Photon Systems Instruments, Czech Republic) (Lee et al., 2021).

### Bacterial growth assay

*X. euvesicatoria* strains (in 10 mM MgCl_2_; OD_600_ 0.0005) were infiltrated into *S. americanum* leaf using needleless syringes. Two leaf discs (1.0 cm^2^) were punched from inoculated leaf area. Leaf discs were ground in 100 μl of 10 mM MgCl_2_. The suspensions were serially diluted and spotted on solid NYG medium containing rifampicin at 20 mg/l. CFU were enumerated 72 h after incubation at 28°C.

### Gene knockout from bacterial genome and complementation

The pK18mobsacB effector-deletion constructs were mobilized into *X. euvesicatoria* by triparental mating following the procedure described by Kvitko and Collmer (2011). Successful deletions were confirmed by PCR using specific deletion-check primers listed in Supplementary Table S1. For complementation, pBBR1:xopOpro:effector-3xFLAG constructs were mobilized into *X. euvesicatoria* D9E strain by triparental mating.

### CRISPR/CasG-mediated gene editing in *S. americanum*

Several specific guide RNAs were designed to target the coding sequences of *SaBs2a* (3), *SaBs2b* (3), *SaBs2c* (3), *SaBs2d* (3) and *SaZAR1* (4) in *S. americanum* SP2273 genome. Total 16 hybridized sgRNAs were assembled into the pDGE vector (Stuttmann et al., 2021) and mobilized into *A. tumefaciens* AGL1 by electroporation. *S. americanum* SP2273 primary transformants were generated by TomatoBioTech (Republic of Korea) and genotyped by sequencing (Macrogen, Republic of Korea) using primers listed in Supplementary Table S1.

### Gene expression analysis

*X. euvesicatoria* strains (in 10 mM MgCl_2_; OD_600_ 0.05) were infiltrated into *S. americanum* leaf using needleless syringes. Four leaf discs (2.0 cm^2^) were harvested from infiltrated leaf area after 28 h. Total RNA was extracted using TRI reagent (Invitrogen, USA) according to the manufacturer’s instructions. Residual genomic DNA was removed by treatment with DNAseI (Sigma-Aldrich, USA). Two μg of RNA was used as template for cDNA synthesis using Maxima First Strand cDNA Synthesis kit (ThermoFisher Scientific, USA). The qRT-PCR analysis was conducted with GoTaq qPCR Master Mix (Promega, USA) using a CFX Connect real-time system (Bio-Rad, USA). The reference gene *Sa18SrRNA* was used for normalization of the samples. Specific primers are listed in Supplementary Table S1.

### Protein extraction and immunoblotting

*A. tumefaciens* AGL1 strains (OD_600_ 0.4) were infiltrated in *N. benthamiana* leaf using needleless syringes. Leaf tissues were frozen after 48 h and ground in liquid nitrogen. Total proteins were extracted in GTEN buffer (10 % glycerol, 50mM Tris-HCl pH 7.5, 2 mM EDTA pH 8, 150 mM NaCl) supplemented with 5 mM DTT, 0.2% IGEPAL (Sigma-Aldrich, USA), 1% PVPP and cOmplete protease inhibitor cocktail (Roche, Germany). Extracts were clarified by centrifugation at 15,000 g for 10 min at 4°C and filtered through MiraCloth (Millipore, USA). Filtered extracts were mixed with 3x SDS sample buffer and denatured at 96°C for 10 min.

*Xe* 85-10 D9E complemented strains were grown in LB medium for 24 h at 28°C. Cells were collected and resuspended in minimal media (0.05% K₂HPO₄, 0.64% KH₂PO₄, 0.1% (NH₄)₂SO₄, 0.035% MgCl₂·6H₂O, 0.01% NaCl and 0.18% fructose) and incubated for 18 h at 28°C to induce expression of type III effector genes. Cells were collected by centrifugation at 3,000 g for 10 min and denatured with 3x SDS loading buffer supplemented with 50 mM DTT.

Proteins were separated by SDS-PAGE and transferred to PVDF membranes. Membranes were probed with anti-FLAG antibodies followed by secondary anti-mouse-horseradish peroxidase antibodies (Sigma-Aldrich, USA). Immunoblots were developed using SuperSignal West substrate (Thermo Scientific, USA) and detected with an Azure 400 CCD imager (Azure Biosystem, USA).

### Statistical analysis

Statistical analyses were conducted with the GraphPad prism 10 software (GraphPad Software, USA) using merged data from at least two independent experiments, variance analysis and appropriate post hoc tests as indicated in figure legends.

## Supporting information

Koh et al_supp info

## Acknowledgements

We thank all members of the Segonzac Lab for their commitment in the initial phase of the project. This work was supported by grants from the National Research Foundation of Korea (NRF) funded by the Korean Ministry of Sciences and ICT (Projects No RS-2025-00512558 and No RS-2024-00349151) to CS and from the Rural Development Administration (RDA; Project No RS-2024-00322297) to CMK.

## Author contributions

Conceptualization: YK, CS; Investigation: YK, HJ, JK, HC, IK, WK; Resources: CMK, KHS, CS; Supervision: KHS, CS; Funding acquisition: CS; Writing: YK, CS with inputs from all the authors.

## Data availability

The data that support the findings of this study are included in this article and in the Supplementary Information.

## Supplementary Information

**Figure S1.** *X. euvesicatoria* strains cause bacterial spot disease symptoms in a commercial tomato cultivar.

**Figure S2.** *Xe* 85-10 effectors trigger cell death upon transient expression in *S. americanum* SP2273.

**Figure S3.** Accumulation of *Xe* 85-10 effector proteins that did not trigger cell death in *S. americanum* SP2273

**Figure S4.** Genotyping of effector deletion in *Xe* 85-10 strains

**Figure S5.** *SaFMO1* expression is reduced in response to *X. euvesicatoria* effector-knockout strains.

**Figure S6.** *SaFMO1* expression is restored in response to D9E(AvrBs2) and D9E(XopJ3) strains.

**Figure S7.** Accumulation of *Xe* 85-10 effector proteins in the D9E complemented strains

**Figure S8.** Identification and expression of *ZAR1* and *Bs2* homologs in *S. americanum*

**Figure S9.** SP2273-*bz* line harbors CRISPR/Cas9-mediated editing at *SaZAR1, SaBs2a*, *SaBs2b*, *SaBs2c* and *SaBs2d* loci.

**Table S1.** Primers used in this study

