## Supplementary material for "*Solanum americanum Bs2* and *ZAR1* homologs recognize *Xanthomonas euvesicatoria* effectors essential for infection": Koh et al_supp info

### Supplementary information

This file contains 9 supplementary figures.

Supplementary Table S1 (primers used in this study) is provided separately.

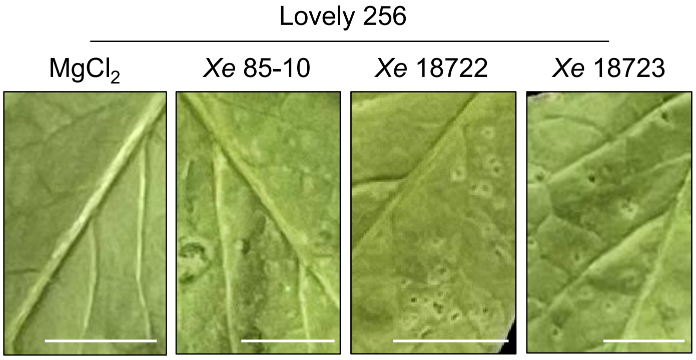


**Figure S1. *X. euvesicatoria* strains cause bacterial spot disease symptoms in a commercial tomato cultivar.**

Lovely 256 plants were dip-inoculated with 10 mM $MgCl_2$, *X. euvesicatoria* strains 85-10 (*Xe* 85-10), 18722 (*Xe* 18722) or 18723 (*Xe* 18723). Leaf area showing disease symptoms were photographed 15 days post-inoculation. Scale bars represent 1 cm.

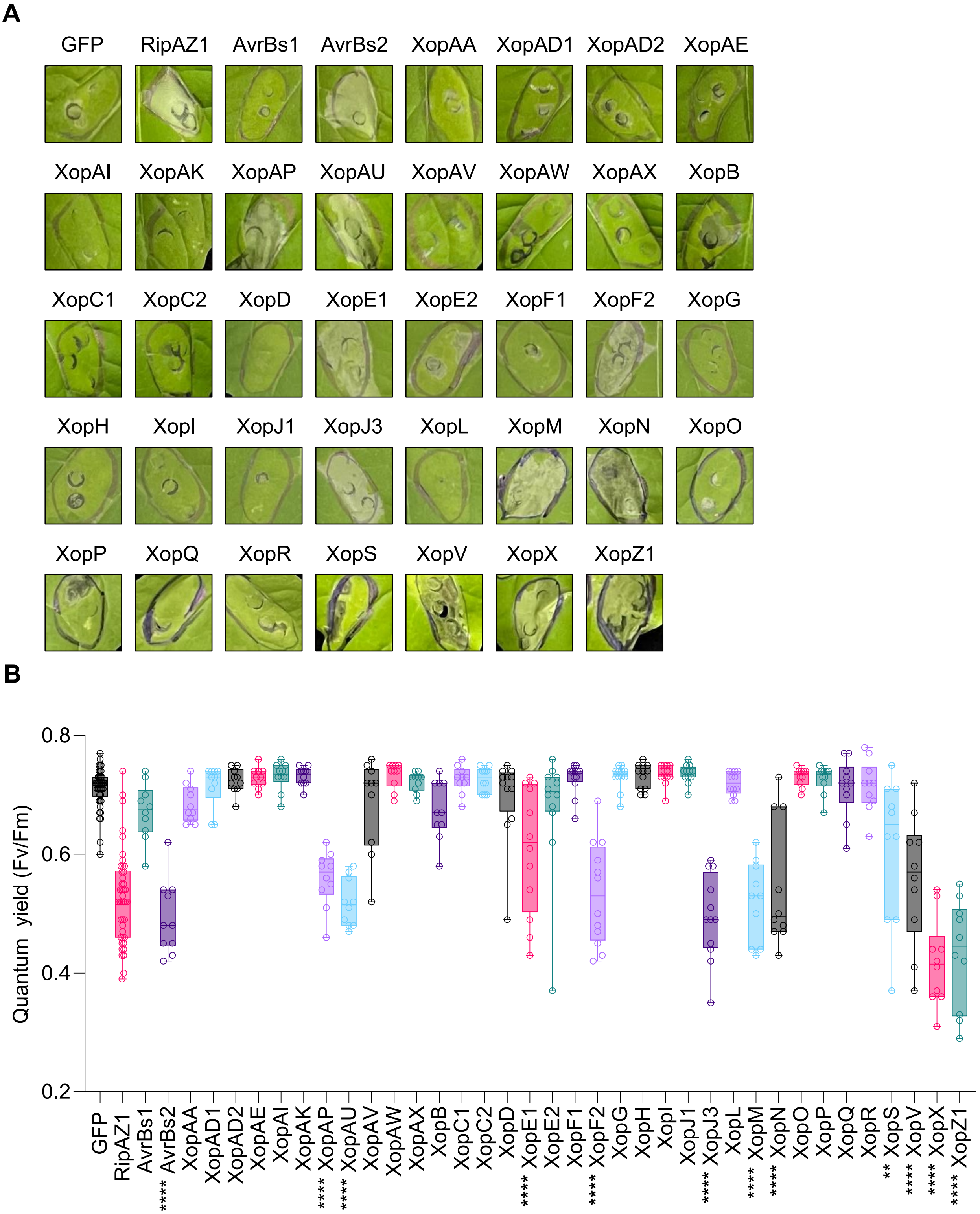


**Figure S2. *X. euvesicatoria* 85-10 effectors trigger cell death upon transient expression in *S. americanum* SP2273.**

**A, B,** Cell death triggered by Agrobacterium-mediated expression of GFP, RipAZ1or each of 37 *Xe* 85-10 effectors in *S. americanum* SP2273. Photographs (bright field) were taken 5 days after infiltration (**A**). Cell death intensity was quantified by measuring the quantum yield (QY) of each infiltrated spot (**B**). High QY indicates strong cell death at the infiltration site, and low QY indicates weak or no cell death. Box plots show the distribution of individual values (n=9 from three biological repeats) between the lower and upper quartiles (25-75%), individual values (dots), and median value (line). The whiskers indicate minimum and maximum values. Asterisks indicate statistical differences with the GFP negative control (one-way ANOVA, Tukey's post-hoc test; **, $P<0.01$; ****, $P<0.0001$).

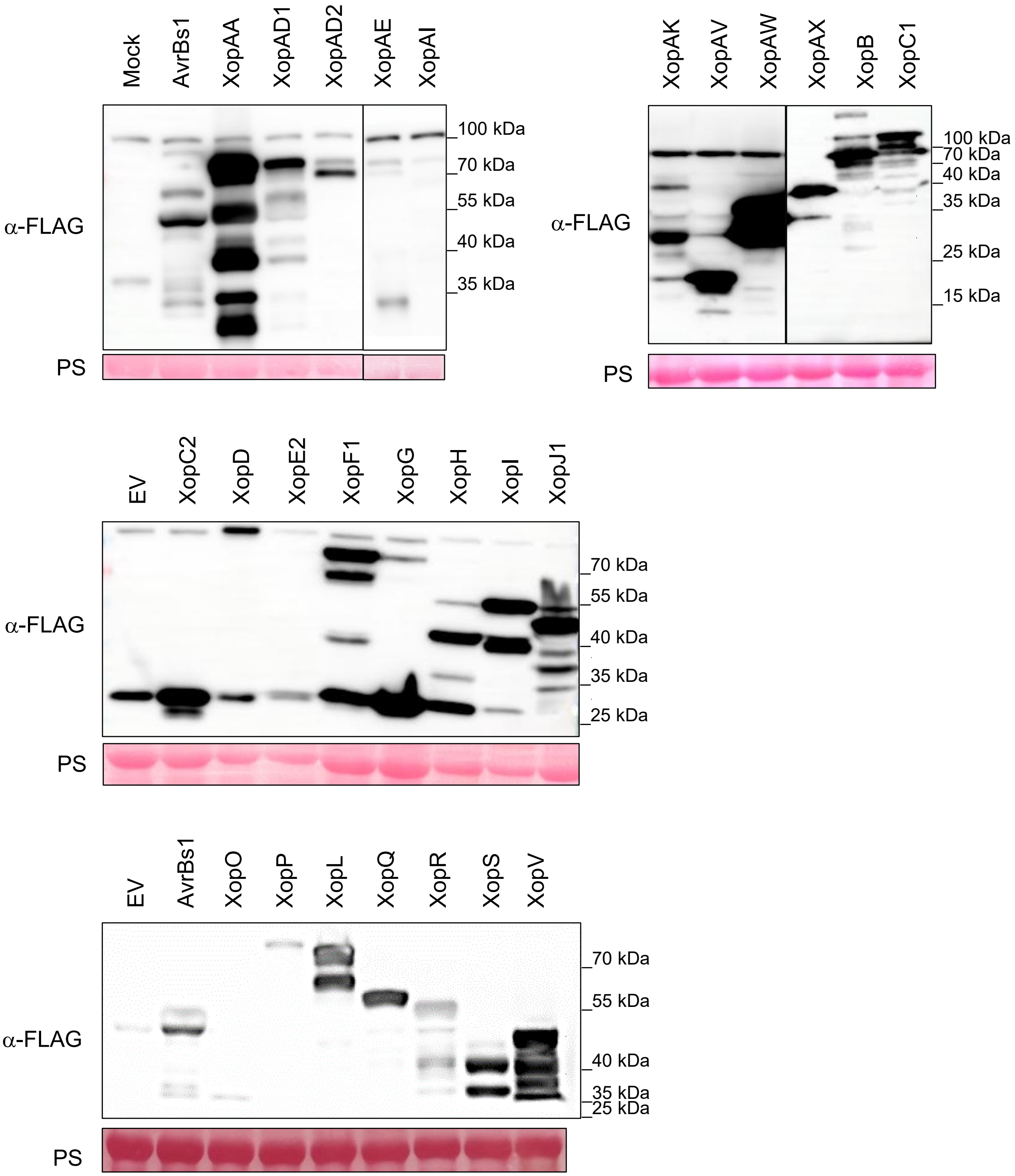


**Figure S3. Accumulation of *X. euvesicatoria* 85-10 effector proteins that did not trigger cell death in *S. americanum* SP2273**

Infiltration buffer (mock) or Agrobacterium strains carrying effector-3xFLAG fusion constructs were expressed in *N. benthamiana*. Leaf samples were harvested 48 h after agroinfiltration. Immunodetection on total protein extracts was performed with anti-FLAG antibodies. Ponceau S staining (PS) shows equal loading of the samples.

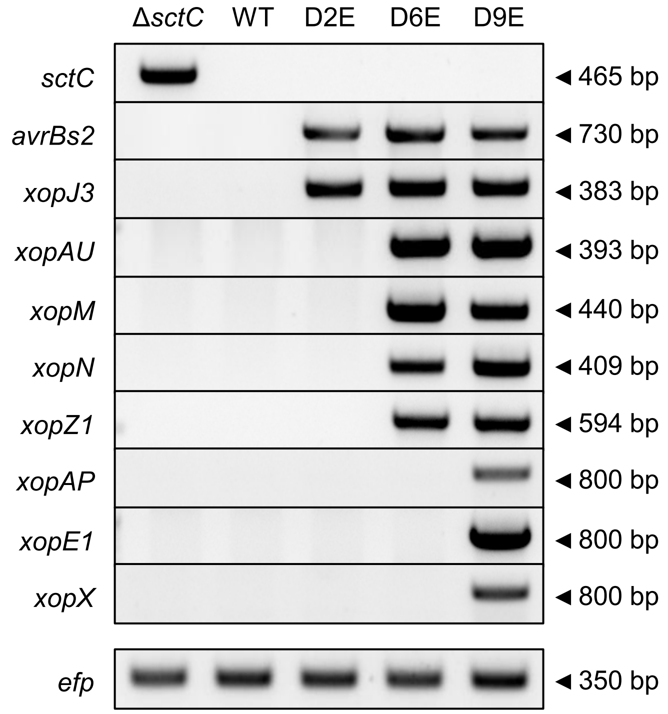


**Figure S4. Genotyping of effector deletion in *X. euvesicatoria* 85-10 strains**

Genomic DNA extracted from *Xe* 85-10 WT, Δ*sctC*, Δ*avrbs2* Δ*xopJ3* (D2E), Δ*avrbs2* Δ*xopJ3* Δ*xopAU* Δ*xopM* Δ*xopN* Δ*xopZ1* (D6E) and Δ*avrbs2* Δ*xopJ3* Δ*xopAU* Δ*xopM* Δ*xopN* Δ*xopZ1* Δ*xopAP* Δ*xopE1* Δ*xopX* (D9E) strains was used as template for amplification across each deleted locus. Amplification of the translation elongation factor P (*efp*) shows equal amount of gDNA template. Specific primers used are listed in Supplementary Table S1. Amplicon length are indicated on the right.

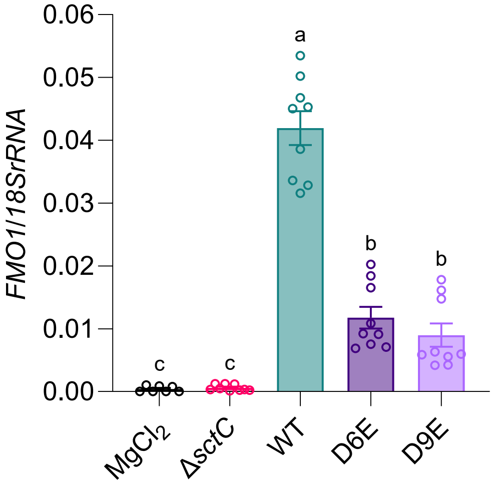


**Figure S5. *SaFMO1* expression is reduced in response to *X. euvesicatoria* effector-knockout strains.**

*S. americanum* SP2273 leaves were infiltrated with 10 mM $MgCl_2$, *Xe* 85-10 WT, Δ*sctC*, D6E or D9E strains. Tissues were harvested 28 h after infiltration for total RNA extraction and qRT-PCR. Dots indicate individual values of *SaFMO1* expression relative to *18SrRNA*; bar graphs represent mean +/- SEM from three biological repeats (n=9). Different letters indicate statistical difference between strains (one-way ANOVA, Tukey's post-hoc test; $P<0.05$).

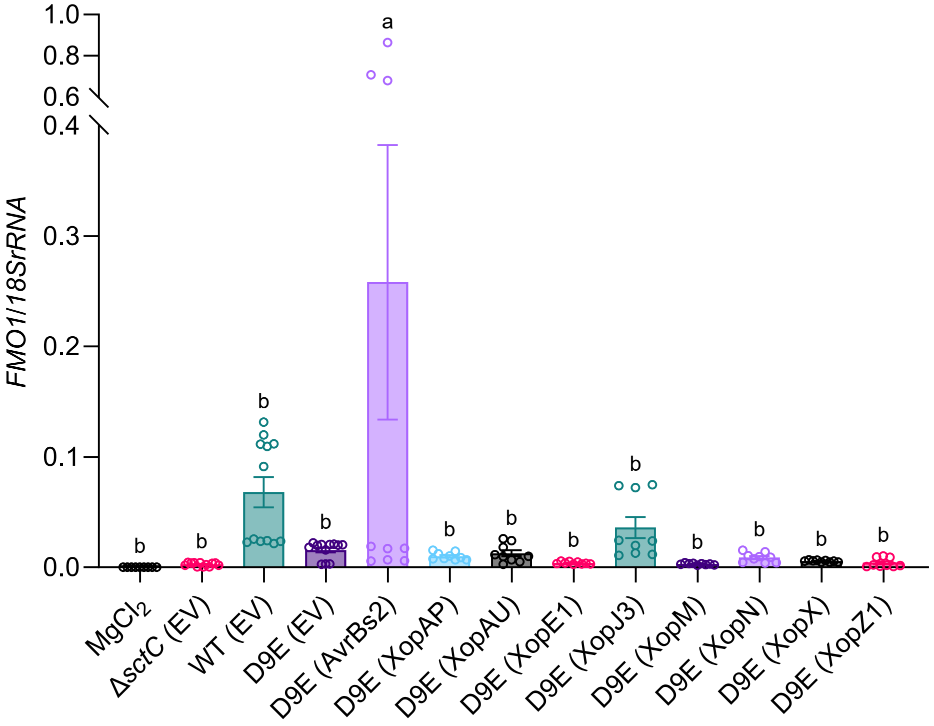


**Figure S6. *SaFMO1* expression is restored in response to D9E(AvrBs2) and D9E(XopJ3) strains.**

*S. americanum* SP2273 leaves were infiltrated with 10 mM $MgCl_2$, *Xe* 85-10 WT, Δ*sctC*, D9E or D9E complemented strains (D9E(EV), D9E(AvrBs2), D9E(XopAP), D9E(XopAU), D9E(XopE1), D9E(XopJ3), D9E(XopM), D9E(XopN), D9E(XopX), and D9E(XopZ1)). Tissues were harvested 28 h after infiltration for total RNA extraction and qRT-PCR. Dots indicate individual values of *SaFMO1* expression relative to *18SrRNA*; bar graphs represent mean +/- SEM from three biological repeats (n=9). Different letters indicate statistical difference between strains (one-way ANOVA, Tukey's post-hoc test; $P<0.05$).

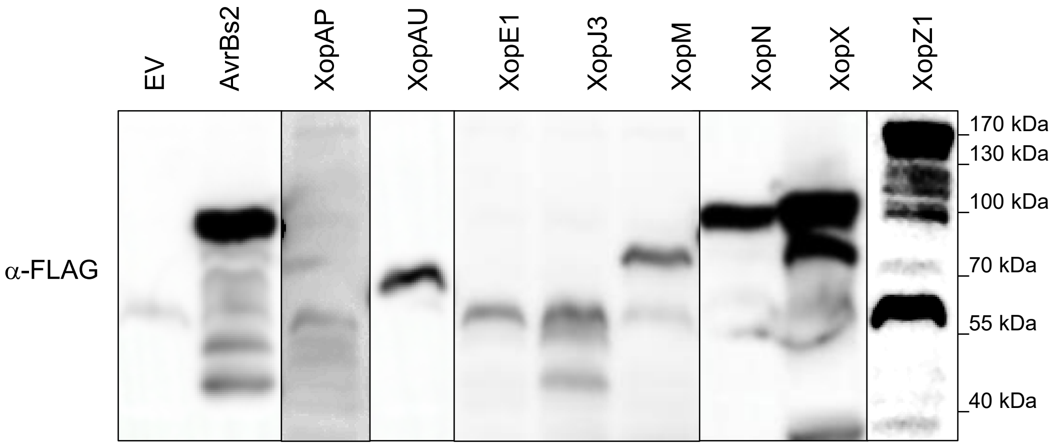


**Figure S7. Accumulation of *X. euvesicatoria* 85-10 effector proteins in the D9E complemented strains**

D9E(EV), D9E(AvrBs2), D9E(XopAP), D9E(XopAU), D9E(XopE1), D9E(XopJ3), D9E(XopM), D9E(XopN), D9E(XopX) and D9E(XopZ1) strains were grown in T3SS-inducing minimal media for 24 h. Immunodetection on total protein extracts was performed with anti-FLAG antibodies.

**A**

| % **Identity** | AtZAR1 | NbZAR1 | SaZAR1 |
|---|---|---|---|
| AtZAR1 | | 57.24 | 58.41 |
| NbZAR1 | 57.24 | | 89.03 |
| SaZAR1 | 58.41 | 89.03 | |

| % **Identity** | Bs2 | SaBs2a | SaBs2d | SaBs2c | SaBs2b |
|---|---|---|---|---|---|
| Bs2 | | 67.19 | 63.90 | 58.13 | 55.54 |
| SaBs2a | 67.19 | | 62.51 | 57.74 | 58.39 |
| SaBs2d | 63.90 | 62.51 | | 59.76 | 59.98 |
| SaBs2c | 58.13 | 57.74 | 59.76 | | 57.89 |
| SaBs2b | 55.54 | 58.39 | 59.98 | 57.89 | |

**B**

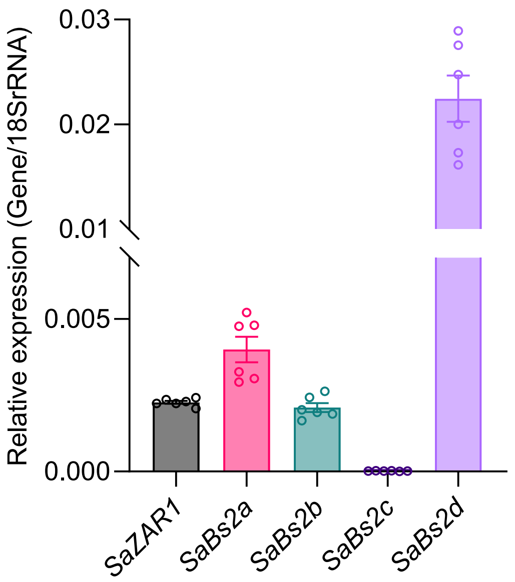


**Figure S8. Identification and expression of ZAR1 and Bs2 homologs in *S. americanum***

**A,** Percentage identity of *S. americanum* proteins homologous to ZAR1 (*Arabidopsis thaliana*, *Nicotiana benthamiana*) or Bs2 (*Capsicum chacoense*). Amino acid sequence alignments were performed with Clustal Omega. **B,** *SaZAR1* and three *SaBs2* homolog genes are expressed in *S. americanum* leaf. *S. americanum* SP2273 leaf tissues were harvested for total RNA extraction and qRT-PCR. Dots indicate individual values of *SaZAR1*, *SaBs2a*, *SaBs2b*, *SaBs2c* and *SaBs2d* expression relative to *18SrRNA*; bar graphs represent mean +/- SEM from three biological repeats (n=9).

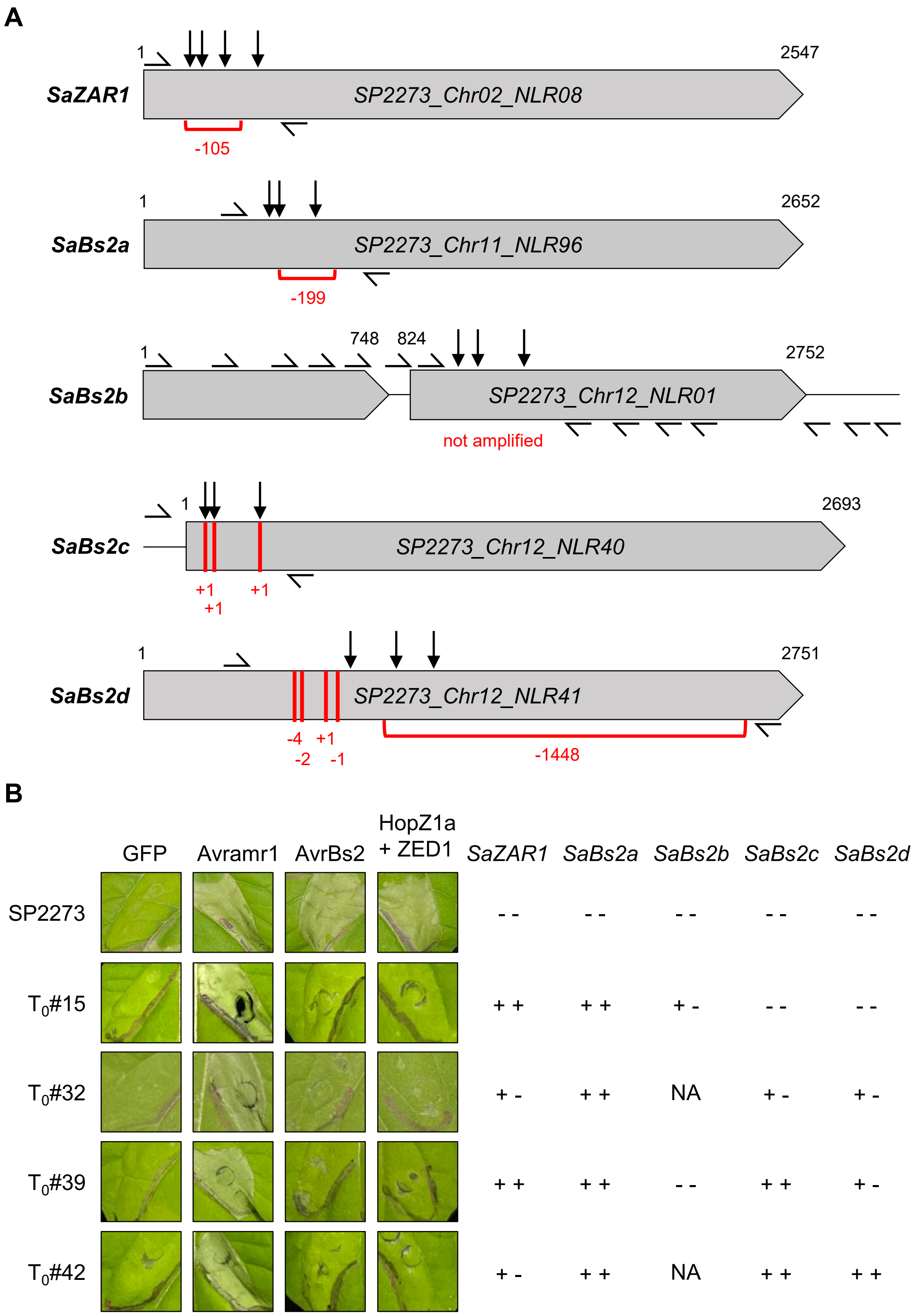


**Figure S9. SP2273-*bz* line harbors CRISPR/Cas9-mediated editing at *SaZAR1, SaBs2a*, *SaBs2b*, *SaBs2c* and *SaBs2d* loci.**

**A,** Gene structure (exons: grey boxes; UTRs, intron: black lines), gRNA position (black arrows), genotyping primers and edits detected in SP2273-*bz* genomic DNA (red lines; + nucleotide insertion; - nucleotide deletion) are indicated for each gene (gene number as annotated in *S. americanum* SP2273 NLRome_CDS by Lin et al., 2023). **B,** Cell death triggered by Agrobacterium-mediated expression of GFP, AvrBs2, HopZ1a + ZED1 or Avramr1 in *S. americanum* SP2273 or SP2273 primary transformants (SP2273-*bz* $T_0$#15, #32, #39, and #42). Photographs were taken 3 days after infiltration. Edition of *SaZAR1*, *SaBs2a*, *SaBs2b*, *SaBs2c* and *SaBs2d* loci is indicated for each transformants (+ +, homozygous mutation; + -, heterozygous mutation; - -, no edition; NA, no amplification).
